# High and low expression of the hyperpolarization activated current (I_h_) in mouse CA1 stratum oriens interneurons

**DOI:** 10.1101/2021.02.26.433114

**Authors:** Lauren T. Hewitt, Gregory J. Ordemann, Darrin H. Brager

## Abstract

Inhibitory interneurons are among the most diverse cell types in the brain; the hippocampus itself contains more than 28 different inhibitory interneurons. Interneurons are typically classified using a combination of physiological, morphological and biochemical observations. One broad separator is action potential firing: low threshold, regular spiking vs. higher threshold, fast spiking. We found that spike frequency adaptation (SFA) was highly heterogeneous in low threshold interneurons in the mouse stratum oriens region of area CA1. Analysis with a k-means clustering algorithm parsed the data set into two distinct clusters based on a constellation of physiological parameters and reliably sorted strong and weak SFA cells into different groups. Interneurons with strong SFA fired fewer action potentials across a range of current inputs and had lower input resistance compared to cells with weak SFA. Strong SFA cells also had higher sag and rebound in response to hyperpolarizing current injections. Morphological analysis shows no difference between the two cell types and the cell types did not segregate along the dorsal-ventral axis of the hippocampus. Strong and weak SFA cells were labeled in hippocampal slices from SST:cre Ai14 mice suggesting both cells express somatostatin. Voltage-clamp recordings showed hyperpolarization activated current I_h_ was significantly larger in strong SFA cells compared to weak SFA cells. We suggest that the strong SFA cell represents a previously uncharacterized type of CA1 stratum oriens interneuron. Due to the combination of physiological parameters of these cells, we will refer to them as <u>L</u>ow <u>T</u>hreshold <u>H</u>igh I_h_ (LTH) cells.

**Key Points:**

- Spike frequency adaptation (SFA) was highly variable among stratum oriens interneurons
- Adapting stratum oriens interneurons were separated into two cell groups using multiple subthreshold and action potential parameters by cluster analysis
- Cells with strong SFA fired fewer action potentials for a given current injection, had lower input resistance and more sag and rebound compared to weak SFA cells
- The physiological differences were not correlated with neuron morphology, location in stratum oriens, or anatomical location along the dorsal-ventral axis of the hippocampus
- Voltage-clamp recordings revealed that strong SFA cells had higher density of the hyperpolarization activated current I_h_ compared to weak SFA cells

## Introduction

The elusive classification of the vast number of GABAergic interneuron subtypes has captivated neuroscientists for decades. The laminar organization of the hippocampus makes it an attractive model for studying circuit mechanisms, particularly because hippocampal interneurons have relatively reliable layer specific inputs and outputs (Cobb et al. 1995; Klausberger and Somogyi 2008; Royer et al. 2012). Hippocampal studies have revealed the extensive diversity of inhibitory interneurons poised to control network timing. Despite comprising merely 10-15% of hippocampal neurons, the remarkably precise activity of GABAergic inhibitory interneurons controls the timing and specificity of excitatory pyramidal cell output and orchestrates layered levels of synchrony in the hippocampal circuit (reviewed in (Bezaire and Soltesz 2013; Freundl and Buzsi 1996; Hájos et al. 2004; Pelkey et al. 2017)). To this end, interneurons play a critical role in facilitating hippocampal activity that ultimately underly complex hippocampal function such as contextual based learning, the formation of episodic memory, and spatial navigation (Lapray et al. 2012; Lovett-Barron et al. 2014; Murray et al. 2011).

Confident classification of the subtype of GABAergic interneurons is necessary in order to understand the particular role an interneuron plays in the local circuitry. Dissection of the functional roles of different inhibitory neurons has been historically difficult. Many studies utilize a three-part approach that entail describing the morphological, biochemical, and physiological properties (Ascoli et al. 2008; DeFelipe et al. 2013). Within the hippocampus, the CA1 region alone contains at least 28 previously described types of inhibitory interneurons located throughout all layers: stratum oriens, stratum pyramidale, stratum radiatum and stratum lacunosum-moleculare (Klausberger and Somogyi 2008). The stratum oriens (SO) of CA1 is home to a variety of regular spiking interneurons, including trilaminar cells, back propagating cells, and oriens-lacunosum moleculare (OLM) cells. OLM cells have a stereotyped morphology with dendrites extending perpendicular to pyramidal cell dendrites along the CA3-subicular axis and an axon that projects through the stratum pyramidale and radiatum until it profusely branches within stratum lacunosum moleculare (Lacaille et al. 1987; Lacaille and Williams 1990; Maccaferri 2005; Maccaferri and McBain 1996). Through this anatomical arrangement, OLM cells coordinate a canonical feedback inhibitory circuit where CA1 neurons activate OLM cells which in turn inhibit the distal dendrites of CA1 pyramidal cells to gate incoming information from the entorhinal cortex via the temporoammonic pathway (Blasco-Ibáñez and Freund 1995; Klausberger 2009; Leão et al. 2012; Maccaferri and McBain 1995; Muller and Remy 2014). OLM cells exhibit tonic or mildly adapting firings characteristics and display sag and rebound responses during the onset and offset of hyperpolarization due to the expression of h-channels (Lupica et al. 2001; Maccaferri and McBain 1996). While OLM cells are well characterized, there are many interneurons throughout the brain discovered in patch-seq (Cadwell et al. 2016; Gouwens et al. 2020), single-cell recording (Tricoire et al. 2011), and single cell transcriptomic (Harris et al. 2018) experiments that remain uncharacterized. Descriptions of the subthreshold and active properties, and the ionic condutances that underly them, in these uncharacterized neurons are paramount to understanding how they contribute to the local circuitry.

In the present study we performed whole-cell recordings from interneurons in the stratum oriens that exhibited oblong cell bodies with dendrites extending along the CA3-subicular axis. We observed heterogenous spike frequency adaptation to depolarizing current injections. Cells with strong spike frequency adaptation (SFA) also had a lower input resistance, larger voltage sag and steeper rebound in response to hyperpolarizing current injections relative to cells with weak SFA. In contrast, single action potential properties were not correlated with SFA. We measured multiple action potential parameters in tandem with subthreshold properties and used a k-means clustering analysis which parsed the cells into two discrete clusters. Using voltage-clamp recordings, we found cells with strong SFA had larger hyperpolarization activated currents (I_h_) compared to cells with weak SFA. We thus suggest the strongly adapting cells are distinct from weak SFA OLM cells. These strong SFA cells exhibit a low-threshold spiking phenotype and will fire action potentials when given small amounts of current relative to other SO interneurons. We will refer to these interneurons as Low-Threshold spiking High I_h_ cells (LTH).

Our data suggests that OLM and LTH cells exhibit distinct firing and subthreshold properties, which are likely dictated from differences in ion channel and biochemical expression profiles. It is a culmination of ion channel expression, inputs and outputs, and intrinsic physiology that will ultimately dictate how a cell responds to different activity states. It is clear that an appreciation for the highly diverse population of inhibitory interneurons is critical to understanding hippocampal function.

## Methods

### Slice Preparation

All animal procedures were approved by the University of Texas at Austin Institutional Animal Care and Use Committee. All mice were housed in a reverse light dark cycle of 12hr on 12hr off with free access to food and water. Experiments used male wild-type (JAX: C57/Bl6, stock #: 000664) and SST:cre Ai14 mice (JAX: SST:cre (stock #:018973) crossed with Ai14 (stock #: 007914)) 2-4 months old (postnatal day 60 – postnatal day 120). Mice were anesthetized with ketamine/xylazine (100/10 mg/kg) and perfused through the heart with ice-cold saline consisting of (in mM): 2.5 KCl, 1.25 NaH_2_PO_4_, 25 NaHCO_3_, 0.5 CaCl_2_, 7 MgCl_2_, 7 dextrose, 205 sucrose, 1.3 ascorbate and 3 sodium pyruvate (bubbled with 95% O_2_/5% CO_2_ to maintain pH at ∼7.4). A vibrating tissue slicer (Vibratome 3000, Vibratome Inc.) was used to make 300 μm thick parasagittal sections from middle hippocampus. Slices were held in a chamber filled with artificial cerebral spinal fluid (aCSF) consisting of (in mM): 125 NaCl, 2.5 KCl, 1.25 NaH_2_PO_4_, 25 NaHCO_3_, 2 CaCl_2_, 2 MgCl_2_, 10 dextrose and 3 sodium pyruvate (bubbled with 95% O_2_/5% CO_2_) for 30 minutes at 35°C and then at room temperature until the time of recording.

### Electrophysiology

Slices were placed in a submerged, heated (33–34°C) recording chamber and continually perfused (1−2 mL/minute) with aCSF 125 NaCl, 2.5 KCl, 1.25 NaH_2_PO_4_, 25 NaHCO_3_, 2 CaCl_2_, 2 MgCl_2_, 10 dextrose and 3 sodium pyruvate (bubbled with 95% O_2_/5% CO_2_). Synaptic transmission was blocked with 20 µM DNQX, 25 µM D-AP5, and 2 µM Gabazine. Slices were visualized with a Zeiss AxioScope or Axioexaminer under 60X magnification. All cells were located in the stratum oriens of CA1 and had oblong cell bodies that ran along the subicular-CA3 axis. Experimenters selected cells with oblong cell bodies, smaller than pyramidal cells, with dendrites extending in the CA3-subcircular axis of the SO, perpendicular to pyramidal cell dendrites. Current injections of 100 pA and -80 pA were delivered to determine if the cells physiological properties were consistent with ‘regular spiking’ adapting cells as described in (Maccaferri and McBain 1996)

### Current Clamp Recordings

Internal recording solution contained (in mM): 135 K-gluconate, 10 HEPES, 7 NaCl, 7 K2 phosphocreatine, 0.3 Na−GTP, 4 Mg−ATP (pH 7.3 with KOH). Current clamp data were acquired using a Dagan BVC−700 amplifier with custom acquisition software written using Igor Pro (Wavemetrics) and sampled at 50 kHz, filtered at 3 kHz and digitized by an ITC-18 (InstruTech) interface. Patch pipettes (4-6 MΩ) were pulled from borosilicate glass. Pipette capacitance was compensated, and the bridge balanced before each recording. Series resistance was monitored throughout each experiment and maintained at approximately 15−35 MΩ. Cells with a series resistance >35 MΩ were omitted from the data set.

### Voltage Clamp Recordings

The internal recording solution was the same as for current clamp recordings (see above). Cells were first recorded in control aCSF (see above) in current clamp mode to record trains of action potentials. Following current clamp recording, voltage gated sodium, potassium and calcium currents were blocked with, 1 µM TTX, 10 mM TEA, 5 mM 4AP, 200 µM BaCl_2_, 100 µM CdCl_2_. Cells were held at -30 mV and inward currents were recorded in response to a series of 1-sec long hyperpolarzing voltage commands (−50 to -130 mV in -10 mV steps). Voltage clamp data was acquired using a Axopatch 200B amplifier (Molecular Devices) with Axograph or custom acquisition software written using Igor Pro (Wavemetrics), digitized at 20 kHz and filtered at 3 kHz. Patch pipettes (4-6 MΩ) were pulled from borosilicate glass.

### Drugs

All drugs (Abcam pharmaceutical or Tocris) were prepared from 1000X stock solutions in water.

### Post-hoc Neuron Visualization

During whole-cell current clamp recordings, neurons were filled with 0.4% neurobiotin. Upon completion of whole-cell recordings, slices were fixed in 3% paraformaldehyde at 4 C overnight. Slices were then washed with 0.1 M PBS 3x for 20 min and place in a blocking buffer of 10% normal goat serum (NGS) and 0.5% triton in PBS overnight at 4 C. Slices were then incubated in PBS containing 1% BSA and 1% NGS with streptavidin conjugated to Alexa-488 (Invitrogen) for 24-48 hr at 4 C. Slices were then rinsed in PBS 3x for 20 min and mounted in Flourmount. Slices were visualized on a resonant scanning 2-photon system (Leica) and Z-stack images were taken. Cells were reconstructed using the max projection of the Z-stack image and analyzed using the SNT plug-in in ImageJ (NIH, https://imagej.net/SNT). (Arshadi et al. 2020)

### K-means clustering

Neuron properties recorded in current clamp (resting membrane potential, steady state R_N_, max R_N_, sag, rebound, ISI ratio, max firing frequency) were used to determine if LTH and OLM cells would separate into distinct populations via k-means clustering. Data was normalized via log transformation and subsequently used for analysis. Using the NbClust package in R (Charrad et al. 2014) we determined the optimal number of clusters for our data set. We then ran a principal components analysis (PCA) and hierarchical clustering on our data and ultimately determined Euclidean distance using k-means analysis.

### Data Analysis and statistical tests

Electrophysiology data were analyzed using custom analysis software written in IgorPro or AxoGraph. The input resistance (R_N_) was calculated from the linear portion of the current-voltage relationship in response to a family of 1500 ms current injections (−40 pA to +40 pA Δ10 pA). The FI curve was calculated from the number of spikes elicited during a family of 1500 ms depolarizing current injections (25 pA to 250 pA Δ25 pA). All statistical analyses (Student’s t-test, ANOVA, repeated measures ANOVA, and Pearson’s correlation) were performed using Prism (Graphpad).

## Results

### Heterogeneity of spike frequency adaptation in low-threshold regular spiking stratum oriens interneurons

We made whole-cell current clamp recordings from cells in stratum oriens of the CA1 region of the hippocampus with oblong cell bodies and horizontally extending dendrites. Low threshold interneurons were identified by having a resting potential close to action potential threshold, an action potential half-width < 1.5ms, and an initial firing frequency < 50 Hz. Stratum oriens interneurons typically display weak to no spike frequency adaptation (Lacaille and Williams 1990; Maccaferri and McBain 1996; Williams and Stuart 2003). Cells were held at -70 mV and depolarizing current injections were used to elicit a train of action potentials. We first measured spike frequency adaptation (SFA) as the ratio of the last interspike interval (ISI) relative to the first ISI (called the ISI ratio) using the first current step that elicited multiple action potentials (Figure 1 A). We found that SFA was highly heterogenous across cells with some showing weak SFA (ISI close to 1) and others showing strong SFA; the ISI ratio was not correlated with the resting membrane potential. (SFA range across all cells: 1.1 – 6.5; Pearson’s r = -0.0015, p = 0.88; Figure 1 B). While ISI ratio is a reliable indicator of SFA, we sought a second method that would reflect SFA across a range of current injections. Using a previously published method for SFA in cortical neurons (Barkai and Hasselmo 1994), we fit a line to a plot of the number of action potentials (spikes) as a function of normalized current injected (Figure 1 C). The area under the linear fit is the S-I value, an indication of adaptation. In agreement with our measurement of ISI ratio, S-I value was highly heterogeneous across all cells recorded (Figure 1 D).

**Figure 1:**
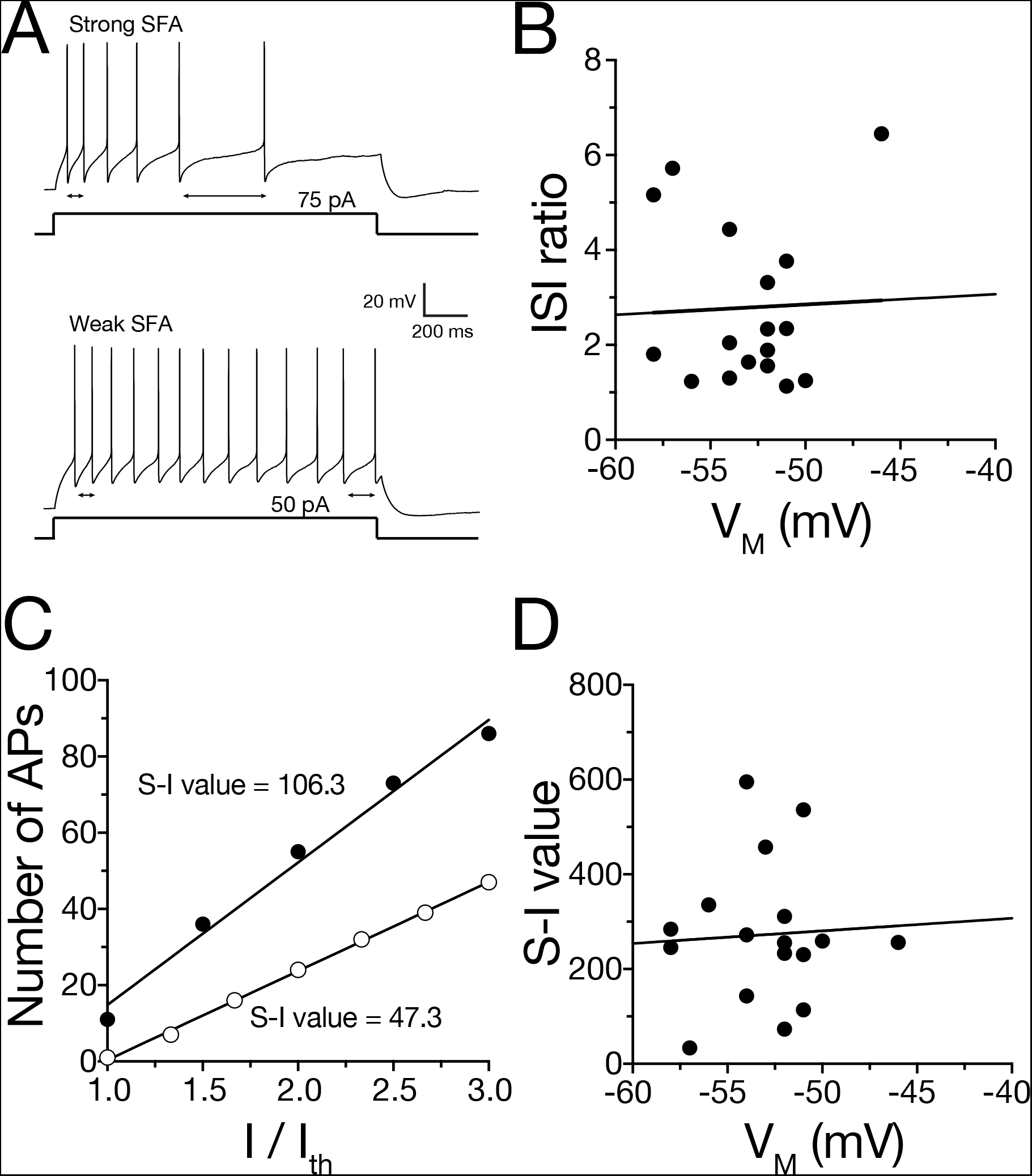
Spike frequency adaptation varies among stratum oriens interneurons A. Voltage traces from weak SFA and strong SFA cells showing the smallest current step to elicit action potentials. Note the strong spike frequency adaptation in the strong SFA cell. Cells were held at -70 mV. B. The ISI ratio does not have a significant relationship to the resting membrane potential (SFA range across all cells: 1.1 – 6.5; Pearson’s r = -0.0015, p = 0.88). ISI ratio was determined by dividing the ISI between the last to spikes (arrow 1) by the ISI of the first two spikes (arrow two). C. Number of action potentials as a function of normalized current injected. The area under the curve (S-I value) indicates the strength of adaptation. D. S-I value is not correlated with VM (mV).

### Relationship of physiological parameters to spike frequency adaptation

The subthreshold properties of hippocampal interneurons, including input resistance (R_N_) and voltage sag (max R_N_ / steady state R_N_), can vary widely across the many interneuron types (for review see (Pelkey et al. 2017)). We measured neuronal input resistance (both maximum and steady state), voltage sag, and rebound and plotted each as a function of SFA (ISI ratio). All four subthreshold parameters were correlated with cell SFA. Maximum R_N_ (Pearson’s r = -0.6047, p = 0.0078) and steady-state R_N_ (Pearson’s r = - 0.6621, p = 0.0028) were negatively correlated with ISI ratio (Figure 2 A-C). Voltage sag (Pearson’s r = 0.7822, p = 0.0002) and rebound slope (Pearson’s r = -0.8941, p < 0.0001) were positively correlated with ISI ratio (Figure 2 D-F; note the negative r value for rebound is because greater rebound is a more negative slope). Like subthreshold properties, action potential properties can vary widely across hippocampal interneurons (for review see (Pelkey et al. 2017)). Unlike subthreshold properties however, there was no correlation between action potential threshold (Pearson’s r = 0.2555; p = 0.3223), action potential half-width (Pearson’s r = 0.1201; p = 0.6462), action potential maximum dV/dt (Pearson’s r = - 0.0637; p = 0.8081), or minimum dV/dt (Pearson’s r = 0.1396; p = 0.5931) (Figure 2 G-L).

**Figure 2:**
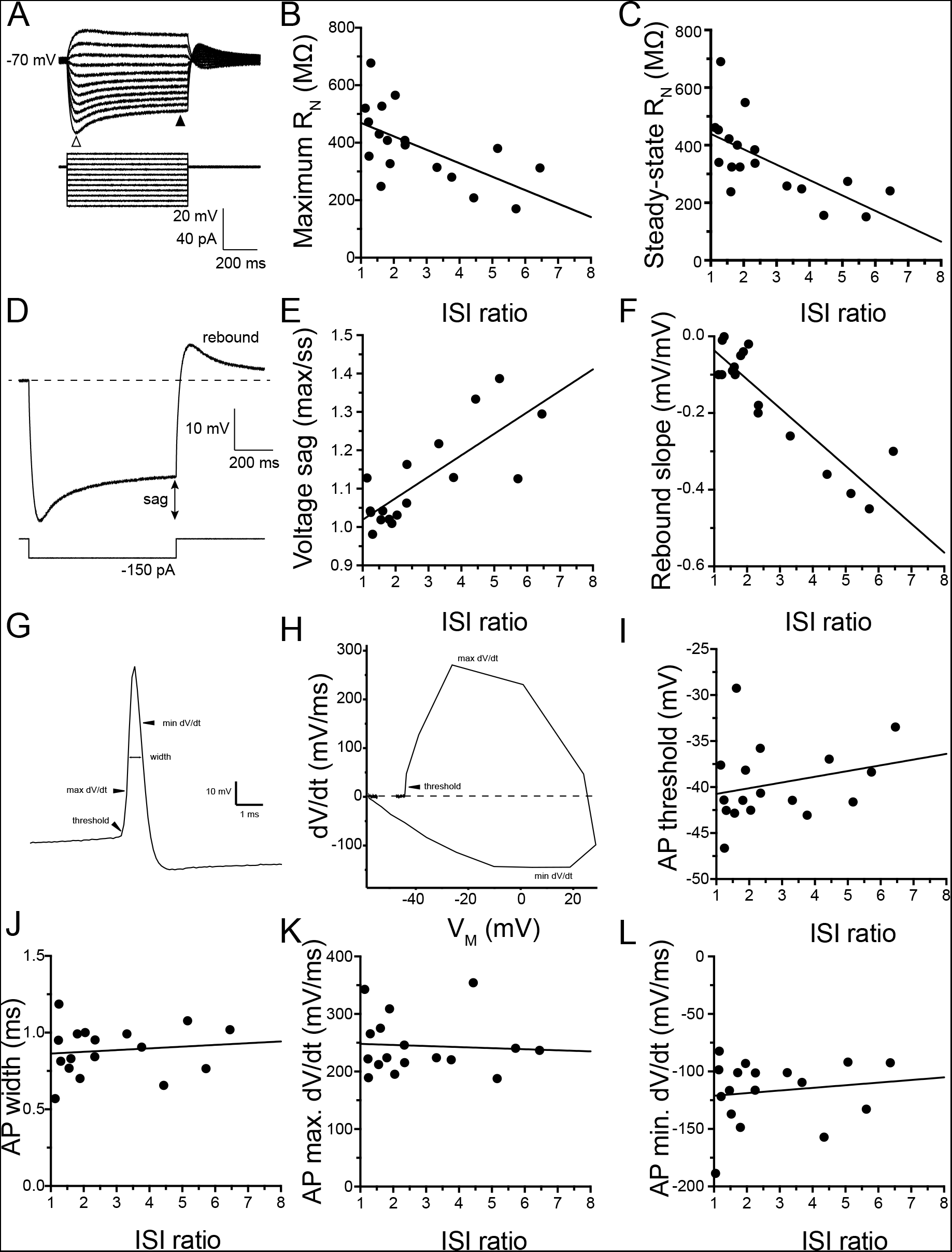
Relationship of physiological parameters to ISI ratio A. Example voltage traces indicating the measurements of maximum (open arrow) and steady state voltages (closed arrow) were taken. B-C. There is a strong correlation of steady state and maximum input resistance with the ISI ratio (Maximum RN (Pearson’s r = -0.6047, p = 0.0078) and steady-state RN (Pearson’s r = -0.6621, p = 0.0028)). D. example voltage traces of the voltage sag and rebound response. E-F. There is a strong relationship between the ISI ratio with voltage sag and rebound (Voltage sag (Pearson’s r = 0.7822, p = 0.0002) and rebound slope (Pearson’s r = -0.8941, p < 0.0001)). G. Example voltage trace of a single AP with arrows indicating where different properties are measured. H. Example phase plane plot of a single AP. Note where maximum dV/dt, minimum dV/dt, and AP threshold are indicated. I-L. Active properties measured from the first action potential. There is no relationship between AP threshold (Pearson’s r = 0.2555; p = 0.3223), half-width (Pearson’s r = 0.1201; p = 0.6462), maximum dV/dt (Pearson’s r = -0.0637; p = 0.8081), or minimum dV/dt (Pearson’s r = 0.1396; p = 0.5931).

### Clustering of heterogenous SFA interneurons

We used these physiological parameters and a k-means clustering algorithm to investigate whether the stratum oriens interneurons we recorded could be separated into functionally distinct categories (Table 1; Figure 3 A). First, we determined the optimal number of clusters given our data set. Our analyses in NbClust in R (Charrad et al. 2014) revealed 2 clusters were optimal (k=2, Figure 3 B). Our data set neatly partitioned into two distinct clusters based off of these physiological characteristics (Figure 3 C-E). Given the large sag and rebound, consistent with the hyperpolarization activated current I_h_, and strong SFA associated with cluster 2, we will tentatively refer to these cells as Low Threshold High I_h_ (LTH) interneurons and refer to cells in cluster 1 as canonical OLM interneurons. We observed OLM (weak SFA) and LTH (strong SFA) neurons with roughly equal probabilities, indicating LTH cells may not be simply a small population of SO interneurons (OLM 10/18 cells, LTH 8/18 cells). Based on the two clusters produced by k-means clustering, we compared the action potential and subthreshold properties between these two groups.

**Figure 3:**
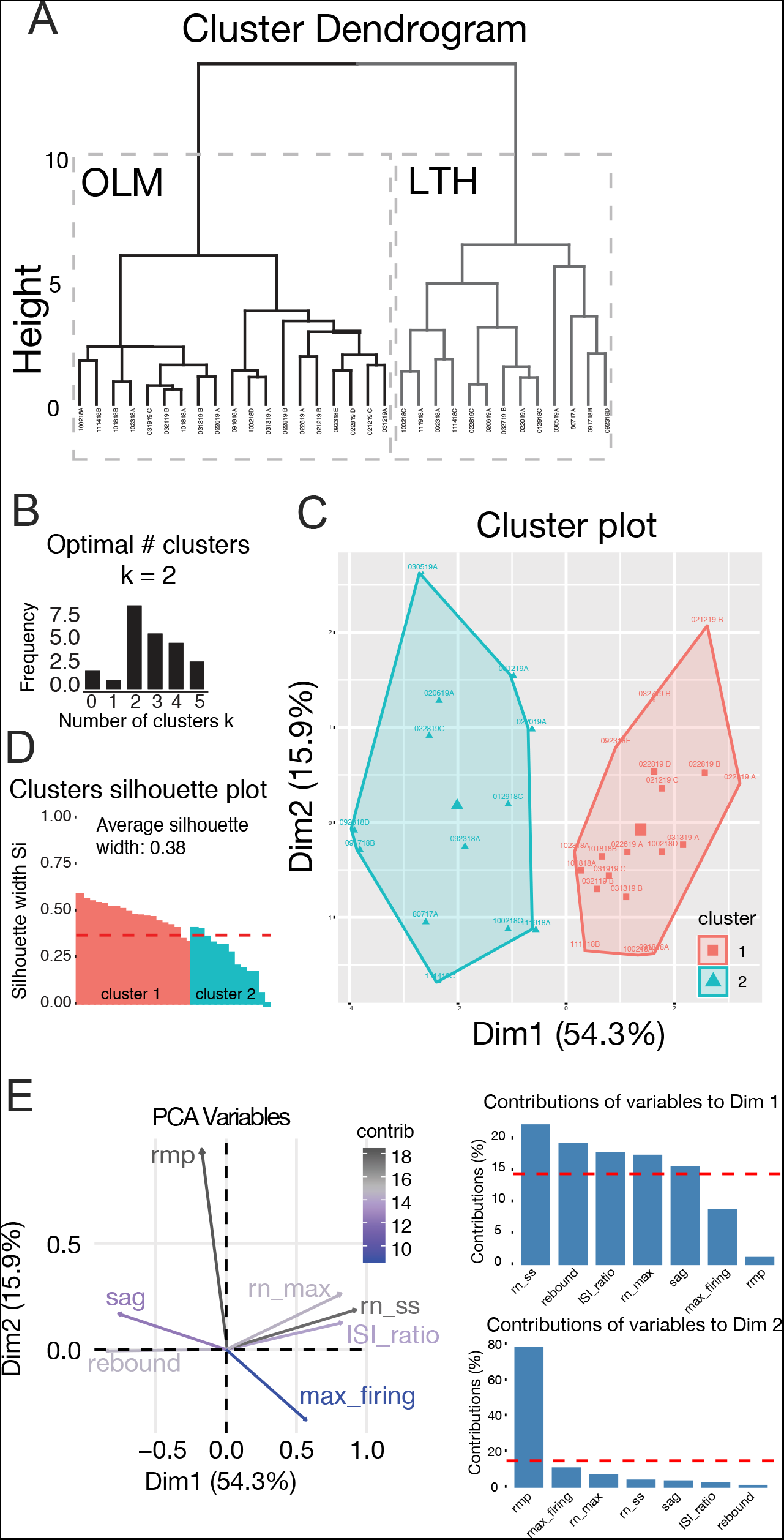
OLM and LTH cells separate into clusters based on intrinsic physiology A. Cluster dendrogram of OLM and LTH cell clusters based on intrinsic electrophysiological properties. B. The optimal number of clusters indicated by NbClust in R Statistics, the optimal number of clusters for our data set k = 2. C. Cluster plot of principal component analysis (PCA) dimensions cluster 1 cells are LTH cells and cluster 2 cells are OLM cells. D. Silhouette plot of clusters 1 and 2. The average silhouette width was 0.38 E. Vector plot of PCA variables and their percent contribution to the clustering of OLM and LTH cells. Bar graphs show the contribution of variability to the two dimensions that carry a majority of the variability of the data set. Bars above the red line indicate variables that contribute more than the calculated average variability of any given parameter.

**Table 1.**
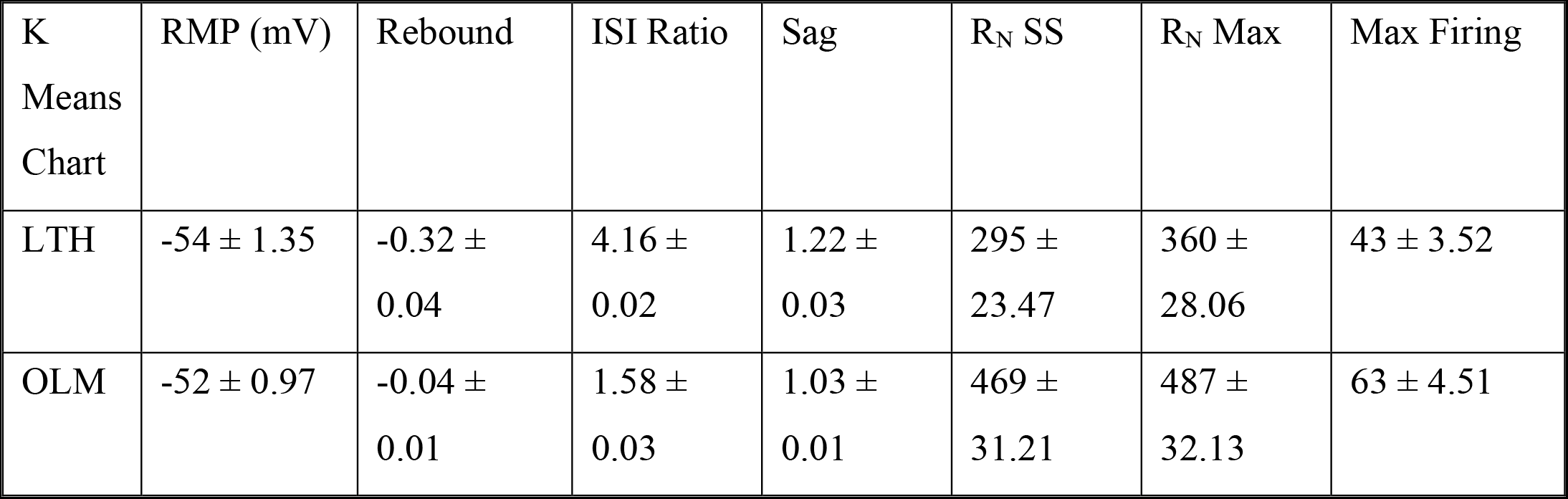
Table of the parameters used in the k-means clustering algorithm from Figure 3. The analysis used the resting membrane potential (RMP, mV), rebound (mV/mV), ISI ratio (first ISI/last ISI), sag (R_N_ SS / R_N_ Max), steady state input resistance (R_N_ SS, Mega Ohm), maximum input resistance (R_N_ Max, Mega Ohm), and max firing rate (Hz).

### Active properties

The results of our clustering analysis suggest our recordings from stratum oriens interneurons can be divided into at least two groups based on physiological characteristics. We next asked if the two groups produced by k-means clustering had different intrinsic properties. We measured action potential firing across a range of depolarizing current injection amplitudes. We found that OLM cells fire significantly more action potentials compared to LTH cells (Figure 4 A-B; repeated measures ANOVA, main effect of cell type (F (9, 144) = 279.3; p = 0.0023)). We isolated the first action potential elicited by the smallest current injection for analysis (Figure 4 C-D). Consistent with our observations in Figure 2, there was no significant difference in action potential threshold, half-width, min or max dV/dt between OLM and LTH cells (Figure 4 E-H). These data suggest the sodium and potassium conductances that contribute to the first action potential were not significantly different between OLM and LTH cells.

**Figure 4:**
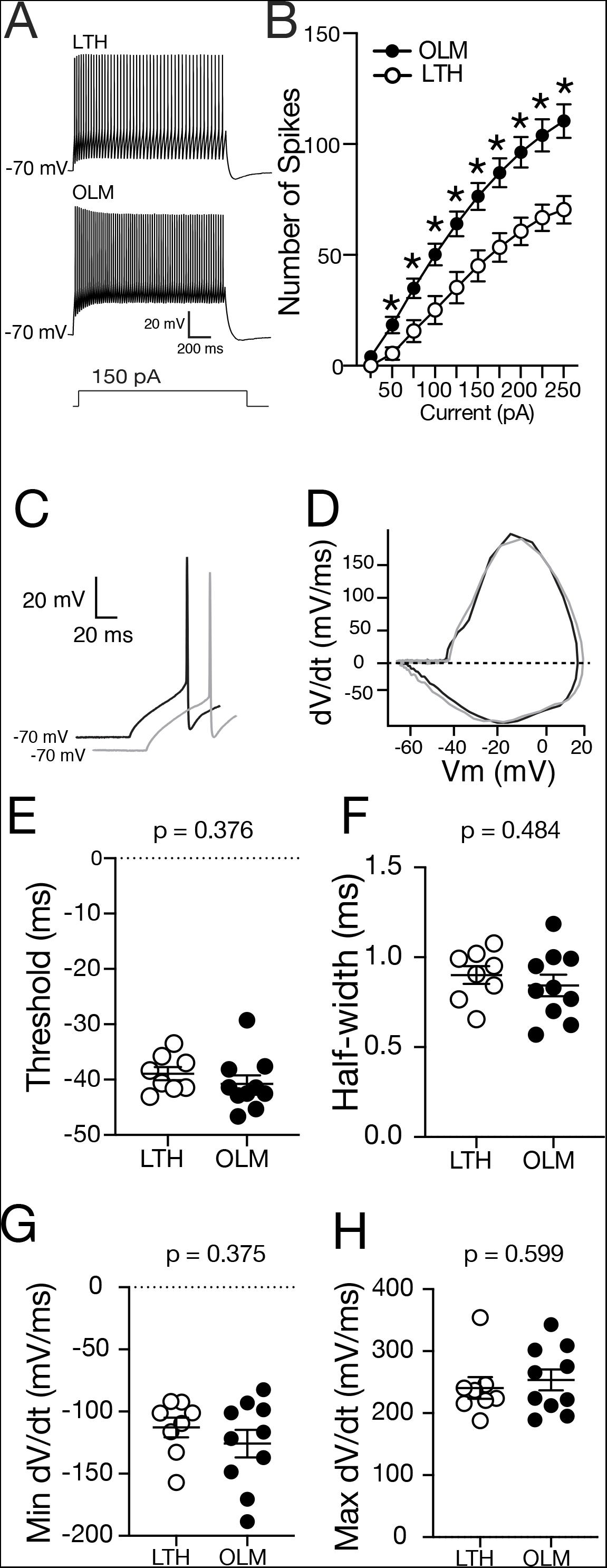
OLM have higher firing rates than LTH cells. A. Voltage traces showing action potential firing from an LTH and OLM cells in response to a 1.5 s, 150 pA current injection. B. Summary plot showing that OLM cells fire significantly more action potentials compared to LTH cells across a range of current injection amplitudes (repeated measures ANOVA, main effect of current (F (9, 144) = 279.3; p = 0.0001), main effect of cell type (F (1, 16) = 13.12; p = 0.0023), interaction between cell type and current (F (9, 44) = 9.66; p = 0.0001). C. First AP on which analysis was conducted. D. Phase plane plot of the first spike. E-H. Action potential threshold (E), half-width (F), minimum dV/dt (G) and maximum dV/dt (H) are not significantly different between LTH and OLM cells.

### Passive Properties

There was no difference in resting membrane potential between OLM and LTH cells (Figure 1 B). In agreement with previous reports on stratum oriens interneurons, both OLM and LTH cells had resting membrane potentials more depolarized compared to hippocampal pyramidal cells (Maccaferri and McBain 1996; Tricoire et al. 2011). OLM cells have significantly higher input resistance (R_N_) compared to LTH cells (Figure 5 B). OLM neurons express h-channels, which is made evident by small voltage sag during the onset and rebound during the offset of hyperpolarizing current injections (Lupica et al. 2001; Maccaferri and McBain 1996; Zemankovics et al. 2010). We measured sag as the difference between the maximum voltage and steady state as indicated by the arrows. (Figure 5 C; grey indicates LTH cells and black indicates OLM cells). While both OLM and LTH cells display sag and rebound, both the sag and rebound were significantly larger in LTH cells compared to OLM cells (Figure 5 D-F).

**Figure 5:**
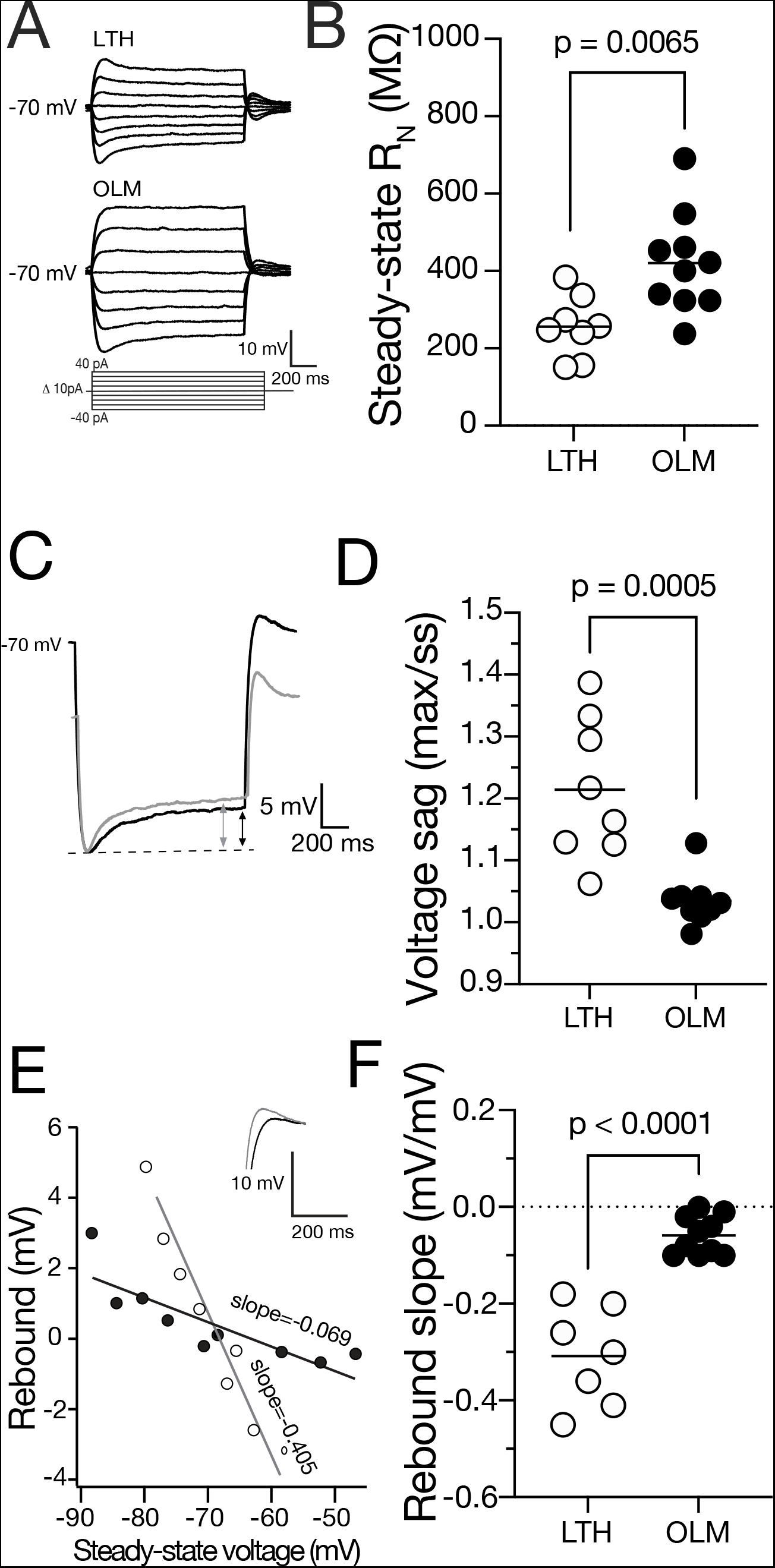
LTH cells have lower input resistance compared to OLM cells. A. Voltage responses to a family of subthreshold current injections from OLM and LTH cells. B. OLM cells have a significantly higher input resistance (RN) than LTH cells (unpaired t-test, p = 0.006). C. Voltage traces indicating the sag in LTH (grey) and OLM (black) cells. The arrows indicate the difference between the maximum voltage and the steady state of the cell. D. LTH cells have significantly higher sag compared to OLM cells (unpaired t-test, p < 0.001). E. Measurement of rebound in the traces seen in C and indicated by the traces of LTH (grey) and OLM (black) cell voltage traces of rebound. F. LTH cells have significantly more rebound than OLM cells (unpaired t-test, p < 0.001).

### OLM and LTH cells express SST

The use of transgenic mouse lines to target interneuron subtypes has been critical to furthering the study of physiological properties of different interneurons (Taniguchi et al. 2011). We recorded fluorescent cells from somatostatin:cre Ai14 mice (SST:cre Ai14) to determine if OLM and LTH cells could both be found in the fluorescent cell population, indicating the expression of somatostatin. We recorded from 13 fluorescent cells (representative images Figure 6 B and D), post hoc analysis of subthreshold and SFA properties indicated 7/13 cells were LTH cells and 6/13 cells were OLM cells. All LTH cells exhibited strong SFA, sag, and rebound properties (Figure 6 A) and had oblong cell bodies in the stratum oriens (Figure 6 B). OLM cells exhibited weak SFA spiking profile with moderate sag (Figure 6 C). Our results suggest the use of SST:cre Ai14 mouse lines indeed lead to sampling a heterogenous population of interneuron subtypes (also indicated in (Hu et al. 2013)), particularly in the stratum oriens where both OLM and LTH cells were found in this mouse model.

**Figure 6:**
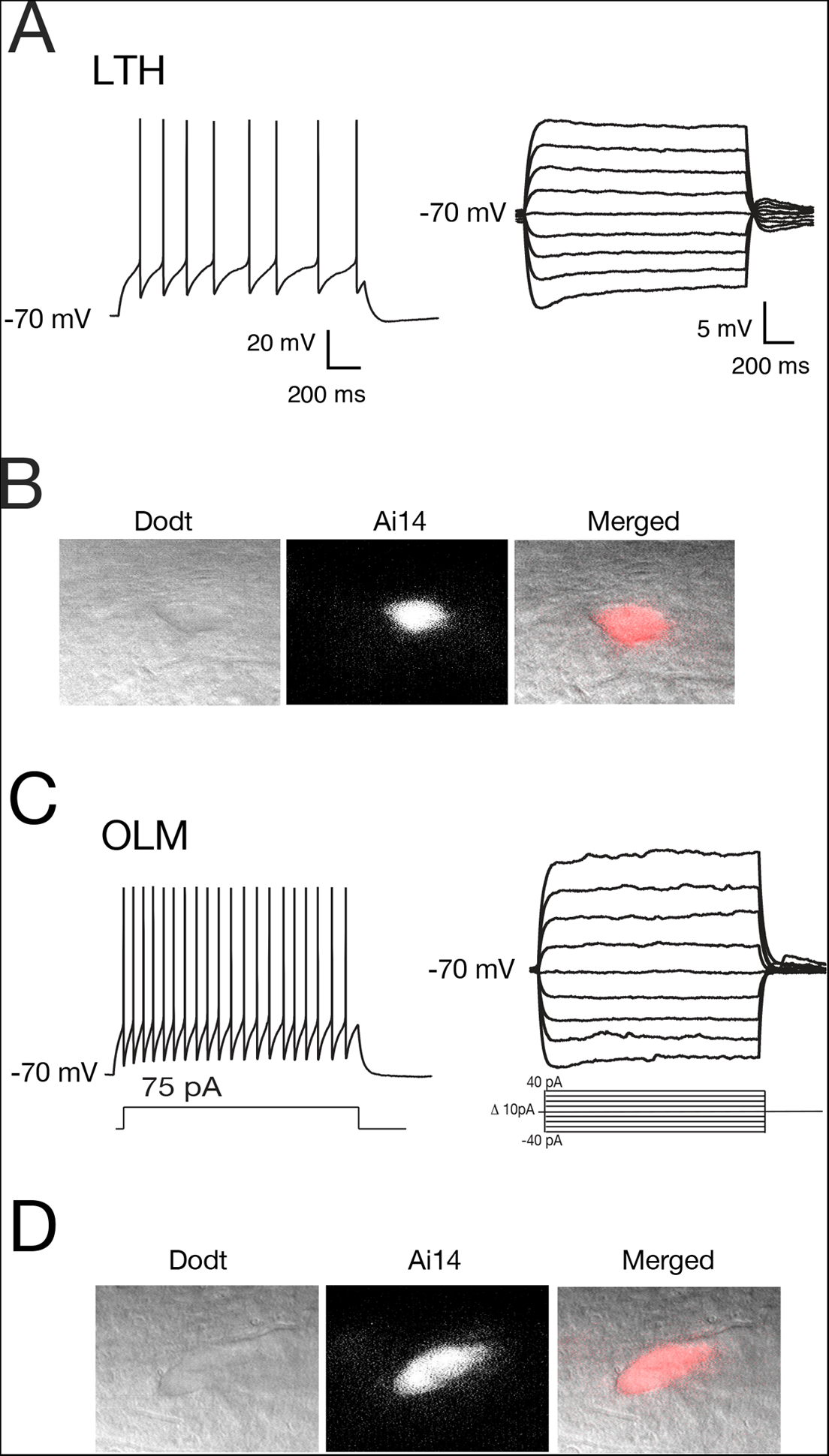
LTH and OLM cells are found in SST:cre Ai14 mice A. representative firing and subthreshold traces of an Ai14+ LTH neuron recording. (n = 7/13 recorded cells) B. LTH interneuron under Dodt contrast, expressing Ai14 driven florescence, and both images overlayed. C. representative firing and subthreshold traces for an Ai14+ OLM neuron recording. (n = 8/13 recorded cells) D. OLM neuron under Dodt contrast, expressing Ai14, and both images overlayed.

### OLM and LTH cells have similar neuronal morphology

The properties of hippocampal pyramidal neurons vary along the dorsal ventral axis of the hippocampus (Dougherty et al. 2012; Kim and Johnston 2015; Malik et al. 2016; Ordemann et al. 2019). More recently, OLM cells were shown to have distinct intrinsic physiological properties depending on the dorsal ventral location of the cell (Hilscher et al. 2019). To determine if the dorsal ventral location of our recordings biased our findings, we mapped our slices to examine their dorsal ventral position within the hippocampus (Malik et al. 2016). Our analysis confirms that our recordings were made from slices taken from the middle hippocampus and that there was no significant difference in the dorsal ventral location of our OLM and LTH cell recordings (Figure 7 A and C). We analyzed the position of our recovered and reconstructed neurons and observed no difference in where OLM and LTH cells are found in the subbicular-CA3 axis of the hippocampus (Figure 7 B). Differences in neuronal morphology will strongly influence physiology. Recorded cells were filled with neurobiotin and reconstructed (Figure 7 D). We did not observe gross differences in the somato-dendritic morphological structure of OLM and LTH cells (Figure 7 E). Both cells had oblong cell bodies in the stratum oriens with dendrites extending along the CA3-subicular axis. A few LTH cells did seem to have dendrites that extended into the pyramidal layer (2 of 6 reconstructions). Axons recovered from OLM cells showed typical extensive branching in the SLM (2 of 5 reconstructions) while axons recovered from LTH cells seemed to also descend out of the pyramidal layer. (2 of 6 reconstructions).

**Figure 7:**
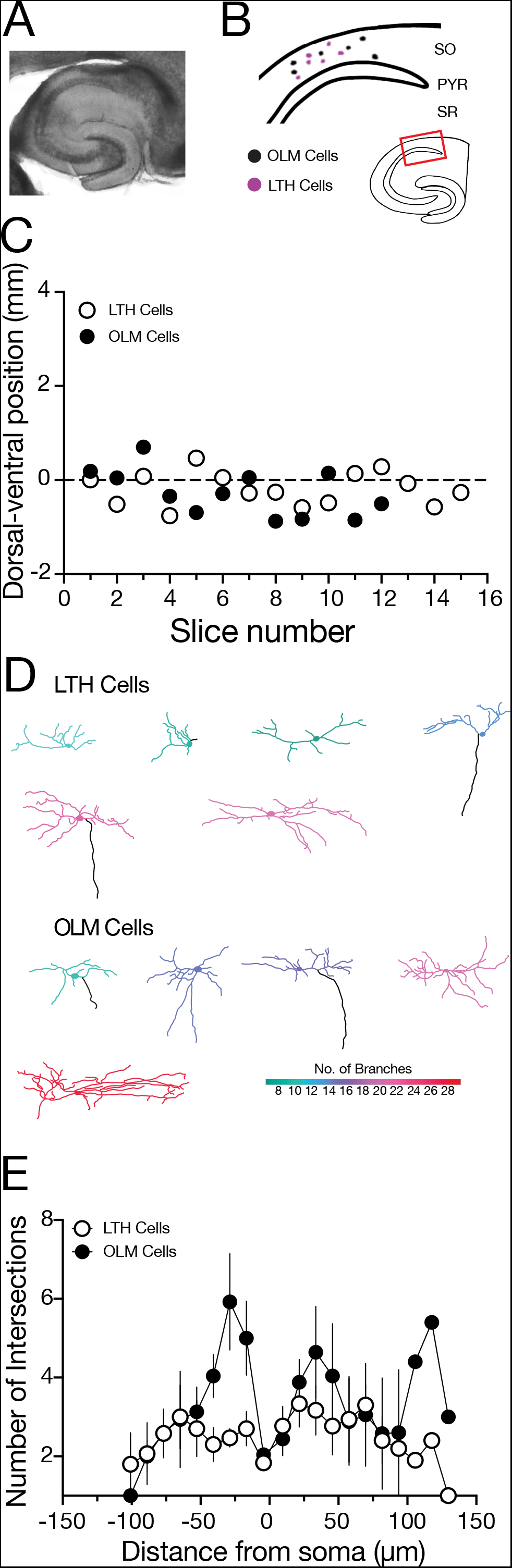
Dorsal/ventral position is not different between LTH and OLM cells, morphology. A. bright field image at 5x showing the CA1 area of the hippocampus. B. schematic of the location of fully recovered and reconstructed LTH and OLM cells recordings in the CA1 SO. Black indicates OLM cell body and pink indicates LTH cell body. C. Dorsal/ventral position of LTH and OLM cells from all slices used in experiments. D. Neuronal reconstructions of LTH and OLM cells. Black lines indicate the axon. The color coding indicates the number of branch points using the scale in the lower right corner. E. Scholl analysis of morphological constructions of LTH and OLM

### Higher I_h_ density in LTH cells

Our current-clamp recordings showed that OLM cells have higher R_N_ and smaller sag and rebound compared to LTH cells. OLM cells were previously shown to have hyperpolarization activated non-selective cation channels (I_h_; (Lupica et al. 2001; Maccaferri and McBain 1996; Matt et al. 2011)). Based on the larger sag and rebound, we hypothesized that LTH cells had higher I_h_ compared to OLM cells. We used a combined current-clamp/voltage-clamp approach to measure the density of I_h_ in OLM and LTH cells (Figure 8 A). After we used depolarizing current injections to evoke action potential firing to measure ISI ratio, we switched to voltage clamp and bath applied 1 µM TTX to block voltage-gated Na^+^ channels; 10 mM TEA, 5 mM 4-AP and 200 µM BaCl_2_ to block voltage-gated K^+^ channels; and 100 µM CdCl_2_ to block voltage-gated calcium channels. In order to compare to previously published results, we used the same whole-cell voltage clamp approach described in Maccafferi and McBain 1996. We used a series of hyperpolarizing voltage steps to elicit a slowly activating inward current (Figure 8 B-C). The density of I_h_ was significantly higher in LTH cells compared to OLM cells (Figure 8, D-E, black circles). The current was completely blocked by the h-channel blocker ZD7288 (50 µM) (Figure 8 D-E, grey circles). These results suggest that the subthreshold differences between OLM and LTH cells were due in part to the differential expression of the hyperpolarization activated current I_h_.

**Figure 8:**
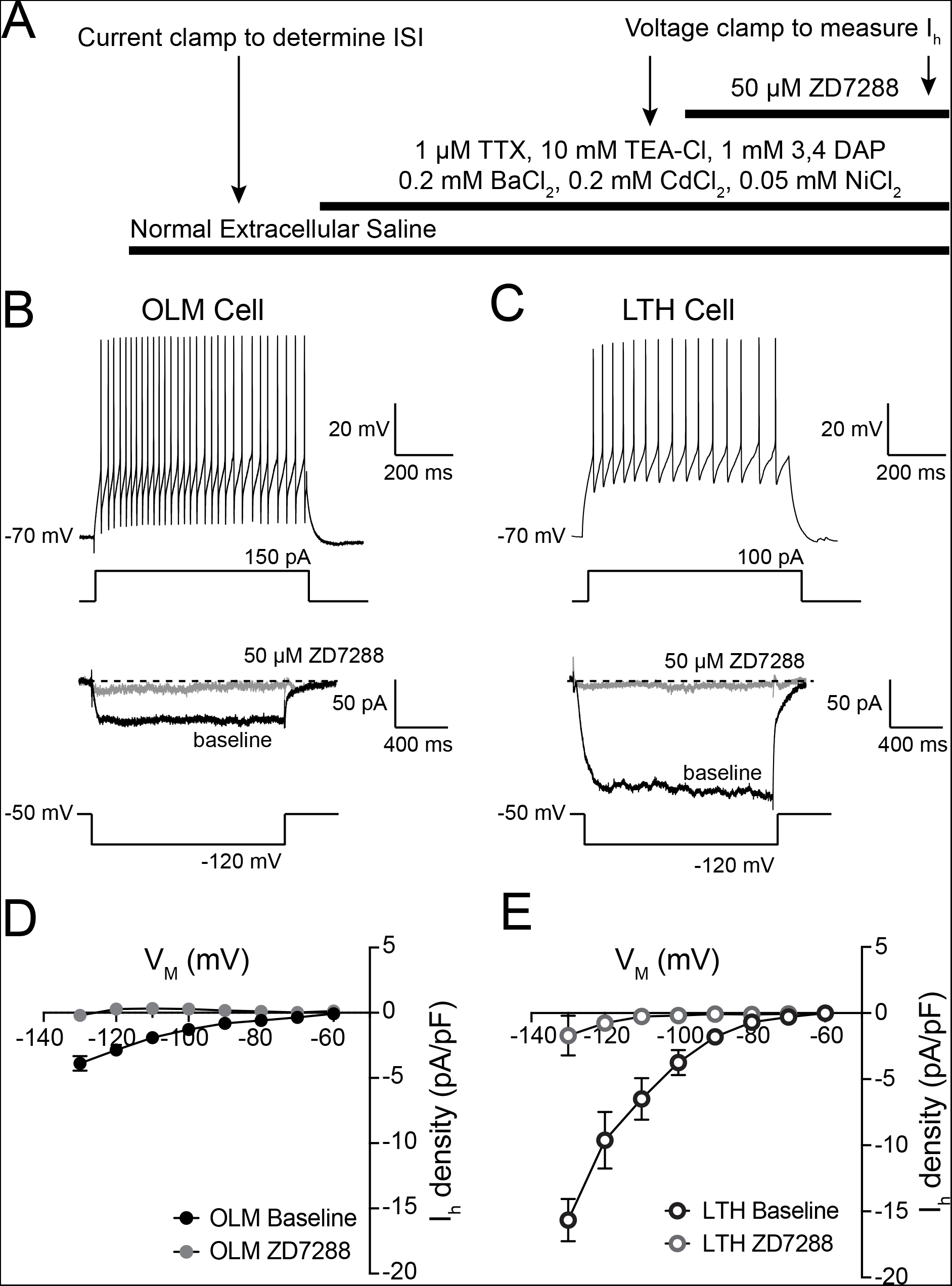
Ih density is higher in LTH cells. A. Experimental design for measuring Ih from OLM and LTH cells. B. Representative OLM cell showing action potential firing with high ISI ratio measured in current clamp (top) and Ih measured in voltage clamp (bottom). C. Representative LTH cell showing action potential firing with low ISI ratio measured in current clamp (top) and Ih measured in voltage clamp (bottom). D. Ih density before and after application of 50 µM ZD7288 in OLM cells. E. Ih density measured before and after application of 50 µM ZD7288in LTH cells.

## Discussion

Interneurons exhibit a wide range of physiological and molecular profiles leading many researchers to question what defines a specific interneuron subtype. Here we use a combination of physiological, morphological, and biochemical approaches to identify a previously uncharacterized subtype of hippocampal stratum oriens interneuron, LTH cells. We identified a number of different intrinsic physiological properties between OLM and LTH cells. LTH cells fire fewer action potentials in response to depolarizing current steps when compared to OLM cells. Single action potential analysis revealed no differences in threshold, amplitude, half-width, or min/max dV/dt, which suggests that the ion channel conductances underlying these properties may not be significantly different between LTH and OLM cells. We found that the input resistance of LTH was significantly lower compared to OLM cells. While the decrease in firing may be related to the lower input resistance of LTH cells, our analysis does not capture how the spike waveform changes with concurrent spikes across the current step, which may reveal an underlying conductance that increases the adaptation of LTH cells leading to a decrease in spikes. Voltage sag was steeper and rebound larger in LTH cells compared to OLM cells. Sag and rebound are current clamp indicators of h-channel function leading us to hypothesize that LTH cells have higher h-channel current compared to OLM neurons. Indeed, we found that I_h_ density is significantly higher in LTH cells compared to OLM cells.

### Ion channel contributions to LTH neuron intrinsic properties

These intrinsic properties of neurons shape the integration of synaptic inputs, precision of action potential output, and contribution to the local circuitry. Our data show LTH cells have significantly higher density of I_h_ resulting in altered subthreshold properties (R_N_, sag, and rebound). Differences in I_h_ between OLM and LTH cells suggest key differences in the ability of LTH neurons to integrate synaptic activity and trigger action potential output. The lower input resistance of LTH cells suggest that larger synaptic currents would be required to fire action potentials. In addition, the higher density of h-channels would also limit the summation of synaptic inputs. Taken together, this suggests that both magnitude and temporal frequency of synaptic inputs would need to be higher to drive LTH cells to fire action potentials compared to OLM cells.

The higher density of I_h_ would be expected to lower input resistance and depolarize the resting membrane potential (Lupica et al. 2001).While we did find that the input resistance of LTH cells was lower compared to OLM cells, there was no difference in resting membrane potential. This could indicate that there is not as much I_h_ at rest in LTH cells or there are likely additional channels active at or near the resting membrane potential may be different between LTH and OLM cells. LTH cells exhibit stronger SFA compared to OLM cells. It is not likely differences in I_h_ density account for the SFA, due to h-channels not being active during large depolarizations. Several different potassium channels can contribute to SFA including Ca^2+^-activated BK and SK channels, and m-channels (Gu et al. 2005, 2007; Peters et al. 2005). Further study is required to determine if there are differences in the expression and or function of these channels between LTH and OLM cells.

### LTH Interneuron Identity

Given the vast diversity of hippocampal interneurons, it is not surprising to find subtypes of interneurons that lack characterization. A previous study dissected different interneuron types based on developmental lineage and uncovered many uncharacterized interneurons (Tricoire et al. 2011). Recent momentous undertakings have revealed that many cell types still lack proper descriptions (Gouwens et al. 2020; Harris et al. 2018). LTH cells appear to be one of these interneuron subtypes that have not been thoroughly characterized. Trilaminar and back propagating interneurons are also found in CA1 SO and present with oblong cells bodies with dendrites that extend perpendicular to pyramidal CA1 neurons (Sik et al. 1994, 1995). It is unlikely that LTH cells are trilaminar cells considering trilaminar cells do not express a sag and rebound response (Gloveli et al. 2005). Back propagating interneurons are typically found in the alveus of CA1, but can also been seen in CA1 SO. These particular interneurons have an axon that extends into CA3. The morphological analysis of the LTH cells in our study did not seem to have any axon processes in CA3, however, this could be due to an incomplete fill of the axon or the axon not being planar in our slice preparation. A paper from Zemankovic et al. in 2010 grouped together cells in the CA1 SO that seemed to be projecting neurons with similar physiological properties and diverse morphological properties (OR group). This group of cells exhibits a steeper sag and rebound response to depolarize steps when compared to OLM cells. However, whole cell voltage clamp study of I_h_ in OR and OLM neurons did not reveal any significant differences (Zemankovics et al. 2010). This could be due to the OR group of neurons being heterogenous indicating LTH cells could potentially be a part of the neurons recorded in the OR group. LTH neurons have very likely been recorded from and included in many studies, here we present a detailed characterization of the intrinsic properties of LTH cells to create a foundation for understanding how these neurons may impact hippocampal circuitry.

### Contrasting LTH and OLM interneurons

It is likely the case, as in many biological systems, that interneuron subtypes cannot be split into discrete subtypes, but instead exist on an axis of properties that define their role in information processing. While many interneuron subtypes do exist on graded axes of properties, given our current data, it seems unlikely for LTH cells to be a subset of the OLM interneurons. While the OLM cells typically originate from the medial ganglionic eminence, a subtype of OLM cells originate from the central ganglionic eminence (Chittajallu et al. 2013; Winterer et al. 2019). These two subclasses of OLM cells are indistinguishable based on physiology. In contrast, we found marked differences in both the firing patterns and subthreshold physiological properties between LTH and OLM cells. OLM interneuron properties can vary based on the dorsal/ventral location of the cell. Ventral OLM cells exhibit more adapting firing rates compared to dorsal cells. Dorsal OLM cells also show a steeper sag response compared to ventral interneurons (Hilscher et al. 2019). We show that both LTH and OLM cells are found in the same dorsal/ventral extent within the middle region of the hippocampus. We further demonstrate a separation of LTH cells from OLM cells based on sampled physiological properties with a k-means clustering algorithm. Taken together, we suggest that LTH cells are a separate class of stratum oriens interneuron.

### Somatostatin expression and morphology of LTH interneurons

While biochemical markers such as parvalbumin (PV) and somatostatin (SST) are traditionally used to differentiate subtypes of hippocampal interneurons, many cells express both PV and SST or other combinations of common inhibitory interneuron markers (Losonczy et al. 2002; Maccaferri et al. 2000; Tricoire et al. 2011). We observed both LTH and OLM interneurons in SST:cre Ai14 leading us to believe LTH cells express SST. Action potential firing is also used to separate classes of interneurons as ‘fast spiking’ vs ‘regular spiking’. Some interneuron classes have distinctive dendritic and axonal arbors but the number of interneurons that can identified by morphology alone, particularly in under characterized interneurons, is limited (Ascoli et al. 2008; DeFelipe et al. 2013; Maccaferri and Lacaille 2003). LTH cells may have an axon that is not planar in our current slice preparation, indeed it would be interesting to investigate the full extent of LTH neuron morphology, particularly the axon, using modern tracing methods. It has become increasingly common to determine the genetic profile of inhibitory interneurons as well, we have gained many insights from recent literature exploring an array of genetic markers of interneuron subtypes that have recorded physiological and reconstructed morphological properties (as in (Gouwens et al. 2020)). Large strides in genetic and big data techniques have allowed scientists to study inhibitory interneurons in exquisite detail. Genetic mouse models and intersectional genetics have made targeted approaches accessible to begin untangling interneuron subtypes functional role in circuits. While we do not show the full molecular profile of LTH cells here, it will be interesting to determine if additional key interneuron markers, if any, are expressed. It is the constellation of properties exhibited by a neuron that determines its function in a circuit. Taking all of these properties into account may be necessary to describe how OLM and LTH cells are both different and related to each other.

## Acknowledgements

We thank Dr. Chris McBain for generously providing protocols for slice processing for reconstructions, assistance with the k-means clustering code, and helpful discussion on this manuscript, Jarod Talbot for conducting the dorsal ventral analysis on processed slices, Dr. Richard Gray for assistance with analysis programs, Meagan Volquardsen for genotyping and mouse colony management, members of the Johnston lab for helpful comments and lively discussion on this manuscript, and to Emily Aster Jones and Jessica Chancey for helpful review of our manuscript. This work was supported by the National Institutes of Health grant RO1 MH100510 (DHB) and the National Science Foundation Graduate Research Fellowship Program (LTH).

## Author Contributions

L.T.H. and D.H.B. designed experiments; L.T.H., G.J.O, and D.H.B. acquired data; L.T.H., G.J.O, and D.H.B. analyzed data; L.T.H. and D.H.B. interpreted results of experiments; L.T.H. and D.H.B. prepared figures; L.T.H. and D.H.B. drafted manuscript; L.T.H., G.J.O., and D.H.B. edited and revised manuscript.

